# Classifying Domains, Benchmarking GPT-4, A Portuguese Dataset for Medical AI Q&A

**DOI:** 10.1101/2024.12.17.627801

**Authors:** Felipe Akio Matsuoka, Henrique Nunes Onaga

## Abstract

Artificial Intelligence (AI), particularly large language models (LLMs), has demonstrated remarkable capabilities in addressing complex tasks, including professional-level medical question answering. While standardized benchmarks like the USMLE have been widely used for evaluating LLM performance in English, there is a significant gap in evaluating these models in other languages, such as Portuguese. To address this, we present a curated dataset derived from the Teste de Progresso (TP), a widely adopted Brazilian progress test used to assess medical knowledge across six key domains: Basic Sciences, Internal Medicine, Surgery, Obstetrics and Gynecology, Public Health, and Pediatrics. The dataset consists of 720 multiple-choice questions spanning five years (2019–2023). We demonstrate two primary applications of this dataset. First, we benchmark the performance of GPT-4, which achieved an overall accuracy of 90% across the six medical domains, with the highest performance in Internal Medicine (10%) and the lowest in Public Health (80%). Second, we develop a classification model based on BERTimbau, achieving an overall accuracy of 94% in categorizing questions into their respective medical domains. Our results highlight the utility of the dataset for both benchmarking AI models and automating medical question classification. This work emphasizes the importance of creating domain-specific datasets in underrepresented languages, like Portuguese, to advance AI-driven medical applications, ensure equitable access to AI technologies, and address linguistic and cultural gaps in healthcare education.

## 1 Introduction

Artificial Intelligence (AI), particularly large language models (LLMs) such as ChatGPT, has demonstrated remarkable capabilities in addressing complex tasks, including answering professional-level exam questions. Standardized tests, such as the United States Medical Licensing Examination (USMLE), have been widely used to benchmark LLM performance, providing structured and objective assessments of cumulative medical knowledge[1, 2]. While these benchmarks have set a strong precedent, they remain predominantly focused on English-language datasets. This limitation poses a challenge for multilingual AI evaluation, particularly in non-English-speaking contexts where the need for robust AI tools in medical education continues to grow.

Brazil, home to one of the largest and most diverse medical education systems globally, provides a compelling case for addressing this gap. The Teste de Progresso (TP)[3], a standardized progress test widely adopted across Brazilian medical schools, offers an ideal framework for benchmarking LLMs on Portuguese-language medical knowledge. Despite the scale and significance of Brazil’s medical education system, there is currently a lack of structured datasets to evaluate AI models in Portuguese, limiting their development and applicability for non-English medical domains.

To address this challenge, we present a curated dataset derived from five years of TP examinations (2019–2023), consisting of 720 multiple-choice questions evenly distributed across six primary medical domains: Basic Sciences, Internal Medicine, Surgery, Obstetrics and Gynecology, Public Health, and Pediatrics. This dataset serves as a valuable resource for benchmarking LLM performance and evaluating their generalization across diverse medical specialties in Portuguese.

While the primary focus of this study is to benchmark LLM performance on Portuguese-language medical knowledge, the inclusion of automated classification demonstrates additional applications of the dataset, further extending its utility. By introducing this resource, we aim to lay the foundation for robust multilingual AI evaluation in medical education and bridge critical gaps in non-English medical knowledge assessment.

This work highlights the importance of structured, domain-specific datasets in fostering advancements in AI-driven medical education. By enabling benchmarking in Portuguese, we contribute to the development of inclusive and equitable AI tools that address the needs of diverse linguistic and cultural settings, particularly within Brazil’s extensive medical education landscape.

## 2 Materials and Methods

### 2.1 Dataset Creation

The methodology of this study focused on two main objectives: the creation of a structured data set from the Teste do Progresso (TP) and the subsequent evaluation of its utility through two experiments. TP is a standardized multiple choice test designed to evaluate the quality of medical education in Brazil. Each test has 120 questions, and they are equally distributed across six main medical areas: Basic Sciences, General Medicine, Surgery, Gynecology and Obstetrics, Public Health, and Pediatrics[4].

Publicly available tests in PDF format were collected from institutional websites on-line. These files were processed to extract and organize the question text, the correct answer, and, where available, the medical topic area associated with each question. PDF files were parsed using Python-based tools such as PyPDF2 and pdfplumber. The text extraction routines were customized to ensure an accurate capture of the question stems and corresponding answer options. The extracted data were then cleaned to remove artifacts introduced during the parsing process, including formatting inconsistencies, incomplete entries, and duplicates. The data set was structured into a CSV file that contains three primary fields: question text, the correct answer, and the question area (when provided).

### 2.2 Subject Area Classification

This study investigated the efficacy of a machine learning model in classifying medical questions based solely on their textual content. To this end, a dataset of medical questions was curated and partitioned into training (70%), validation (15%), and test (15%) subsets, ensuring a balanced representation of questions across all medical subject areas.

A BERTimbau model[5], pre-trained on Brazilian Portuguese text, was employed as the foundational architecture. This choice was motivated by BERTimbau’s demonstrated efficacy in capturing semantic relationships within Portuguese text, aligning with previous research that successfully leveraged this architecture for similar regression tasks in the Portuguese language[6]. An output layer, tailored to categorize questions into six distinct medical domains, was appended to the BERTimbau architecture.

The model was trained using the AdamW optimizer[7], a variant of stochastic gradient descent that incorporates weight decay and momentum, with a learning rate of 2e-5. A linear learning rate schedule, devoid of a warmup period, was employed, ensuring a consistent learning rate throughout the training process. To mitigate the risk of exploding gradients, a phenomenon known to destabilize training, gradient clipping with a maximum norm of 1.0 was applied. Model performance was evaluated over 4 epochs, with each epoch comprising iterative training data batch processing, cross-entropy loss computation, and backpropagation for parameter updates.

All experiments were conducted utilizing the computational resources of Google Colab, specifically leveraging a T4 GPU with 15GB of VRAM.

### 2.3 GPT-4 Perfomance Analysis

The second experiment assessed the performance of the ChatGPT-4 language model in answering multiple-choice questions. The Interinstitutional Progress Test was chosen as the evaluation benchmark, consisting of 120 multiple-choice questions. Each question was input into a new chat session using a zero-shot standardized prompt.

The model’s responses were systematically recorded and compared against the correct answers to determine its accuracy. Accuracy served as the primary evaluation metric, providing a quantitative measure of the model’s performance. To gain further insights, an error analysis was conducted, focusing on patterns in the model’s incorrect predictions. This analysis examined whether specific subject areas or question types posed greater challenges for the model, shedding light on potential limitations in its capabilities.posed greater difficulty.

#### Prompt

“I will provide you with a multiple-choice question with only one correct answer. Return the correct alternative.”

## 3 Results

After cleaning and validation, the final dataset contained a total of 720 questions, with 240 questions annotated with the corresponding medical subject area.

### 3.1 Subject Area Classification

The model demonstrated promising performance on the held-out test set, achieving an overall accuracy of 94%. This indicates that the model correctly classified 94% of the medical questions in the test set.

A detailed examination of the per-class performance reveals consistently high precision and recall across all six medical subject areas. Notably, Basic Sciences and Public Health achieved perfect precision and recall (1.00), signifying flawless classification in these categories. General Medicine exhibited slightly lower precision (0.83) but maintained perfect recall (1.00), suggesting that while all instances of this class were correctly identified, some instances from other classes were misclassified as General Medicine. Pediatrics and Surgery also demonstrated strong performance, with F1-scores of 0.92 and 0.93, respectively, indicating a good balance between precision and recall. Gynecology and Obstetrics also performed well, further validating the dataset’s robustness in classifying diverse medical domains (Table 2).

**Table 1:** Performance of ChatGPT across six medical domains based on the Interinstitutional Progress Test.

| Medical Domain | Correct Answers | Total Questions | Accuracy |
| --- | --- | --- | --- |
| Basic Sciences | 17 | 20 | 85% |
| Internal Medicine | 20 | 20 | 100% |
| Surgery | 18 | 20 | 90% |
| Obstetrics and Gynecology | 19 | 20 | 95% |
| Public Health | 16 | 20 | 80% |
| Pediatrics | 17 | 20 | 85% |
| **Overall Performance** | **108** | **120** | **90%** |

**Table 2:**
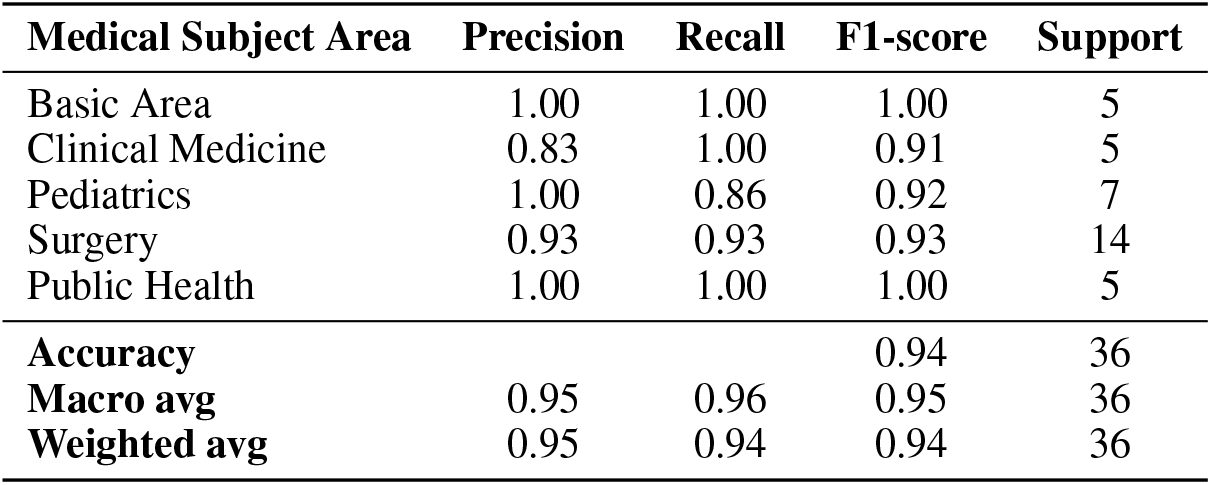
Model Performance on Medical Question Classification.

### 3.2 GPT-4 Perfomance

ChatGPT-4 demonstrated strong performance in answering multiple-choice questions from the Interinstitutional Progress Test, achieving an overall accuracy of 90%, correctly answering 108 out of 120 questions. Performance varied across the six medical domains, with the highest accuracy observed in Internal Medicine, where the model answered all 20 questions correctly, achieving score of 100%. Obstetrics and Gynecology followed with 95% accuracy (19/20 correct answers), while Surgery demonstrated a 90% accuracy rate (18/20). In the Basic Sciences and Pediatrics domains, the model achieved 85% accuracy, correctly answering 17 out of 20 questions in each. The lowest performance was observed in the Public Health domain, with 16 correct answers out of 20, corresponding to an accuracy of 80%.

## 4 Discussion

The findings from our study underscore the utility of the dataset in evaluating both domain-specific question classification and the performance of advanced language models like ChatGPT-4. The dataset, carefully curated to cover diverse medical domains, demonstrated its effectiveness in benchmarking models and assessing their ability to process and classify medical questions accurately. The BERTimbau-based model achieved high accuracy and strong per-class performance, highlighting its potential for medical text applications, such as automated medical question routing and preliminary diagnostic assistance based on textual content. These results reinforce the value of the dataset in developing and evaluating machine learning models for healthcare applications.

For the ChatGPT-4 experiment, the dataset provided a robust framework for assessing the model’s strengths and limitations across medical domains. ChatGPT-4 achieved strong accuracy overall. However, errors were distributed across specific areas. In Obstetrics and Gynecology, mistakes often stemmed from misinterpretation of nuanced clinical scenarios, while in Public Health, errors were related to statistical interpretation and policy-focused questions. In Basic Sciences and Pediatrics, errors appeared sporadically, mainly in questions involving detailed physiological or clinical processes. These patterns of errors reveal the dataset’s ability to probe the contextual and domain-specific understanding of language models, emphasizing its relevance for benchmarking AI systems.

Despite these promising results, the current study is not without limitations. The relatively small size of the test set may constrain the generalizability of the findings. Future work should focus on expanding the dataset to include a larger and more diverse set of questions, covering additional medical domains and subdomains. Such efforts would not only enhance the dataset’s representativeness but also improve its utility for evaluating the robustness and scalability of both domain-specific models and general-purpose language models.

Furthermore, this study highlights the critical importance of developing and utilizing datasets in Portuguese, particularly for medical question-and-answer tasks. With the growing interest in AI applications tailored to specific languages and regions, having datasets like ours ensures that models can be effectively benchmarked and applied in real-world contexts. By addressing the need for high-quality datasets in Portuguese, our work paves the way for advancing AI solutions that cater to linguistic and cultural nuances in healthcare.

In conclusion, our results emphasize the dataset’s versatility and potential for fostering advancements in AI for medical applications. It serves as a valuable tool for benchmarking models, understanding their strengths and weaknesses across medical domains, and driving further research into automated systems for medical education and decision support. The importance of datasets in underrepresented languages, such as Portuguese, cannot be overstated, as they play a vital role in ensuring equitable access to cutting-edge AI technologies in diverse healthcare settings.

## 5 Data Availablity

The dataset used in this study, along with the code for the classification model, will be made publicly available on GitHub upon publication of the paper. This will include the full dataset, detailed documentation, and the implementation of the classification model to ensure transparency and facilitate reproducibility of our findings. The GitHub repository link will be provided in the final version of the paper.

## References

[1] Tim H. Kung, Matthew Cheatham, Annabelle Medenilla, Charmaine Sillos, Lorena De Leon, Clarence Elepaño, Mark Madriaga, Ryan Aggabao, Gerardo Diaz-Candido, Joshua Maningo, and Victor Tseng. Performance of chatgpt on usmle: Potential for ai-assisted medical education using large language models. PLOS Digital Health, 2(2):e0000198, 2023.

[2] Hanjie Chen, Zhouxiang Fang, Yash Singla, and Mark Dredze. Benchmarking large language models on answering and explaining challenging medical questions, 2024.

[3] Angélica Maria Bicudo, Pedro Tadao Hamamoto Filho, Joelcio Francisco Abbade, Maria de Lourdes Marmorato Botta Hafner, and Cláudia Maria Leite Maffei. Teste de progresso em consórcios para todas as escolas médicas do brasil. Revista Brasileira de Educação Médica, 43(4):151–156, 2019.

[4] Associação Brasileira de Educação Médica. Teste de progresso em consórcios para todas as escolas médicas do brasil. Revista Brasileira de Educação Médica, 43(4):191–200, 2019.

[5] Felipe Souza, Rodrigo Nogueira, and Roberto Lotufo. Bertimbau: Pretrained bert models for brazilian portuguese. In Intelligent Systems: 9th Brazilian Conference, BRACIS 2020, Rio Grande, Brazil, October 20–23, 2020, Proceedings, Part I, pages 403–417. Springer International Publishing, 2020.

[6] Felipe Akio Matsuoka. Automatic essay scoring in a brazilian scenario, 2023.

[7] Ilya Loshchilov and Frank Hutter. Decoupled weight decay regularization. In International Conference on Learning Representations, 2019.

